# Infraslow modulation of theta synchrony in the hippocampus circuit during REM sleep

**DOI:** 10.64898/2026.01.30.702893

**Authors:** Nelson Espinosa, Mauricio Caneo, Guillermo Lazcano, Alejandro Aguilera, Ariel Lara-Vasquez, Pablo Fuentealba

## Abstract

Sleep dynamically reorganizes hippocampal activity, but how this reconfiguration supports large-scale coordination across the hippocampal circuit remains unclear. We performed simultaneous recordings from dorsal (CA1d) and ventral (CA1v) hippocampus, retrosplenial cortex (RSC), prefrontal cortex (PFC), and dorsal thalamus in rats across brain states. Theta oscillations dominated hippocampal activity during active behaviour and REM sleep, while delta oscillations prevailed during nREM and quiet waking. Phase synchrony between CA1d and CA1v varied across states, peaking during REM sleep and exhibiting infraslow fluctuations (0.02 Hz) unique to this stage. These slow modulations persisted after controlling for theta power, suggesting a genuine modulation of interregional synchrony. Transitions into REM were marked by rising CA1d–CA1v synchrony and widespread neuronal activation, with thalamic activity most strongly predicting coupling dynamics. During REM sleep, neuronal firing across hippocampal–cortical–thalamic circuits was phase-locked to the infraslow theta coupling cycle, indicating gain modulation at the network level. Such entrainment was strongest and most consistent in REM and aligned near a common phase, revealing temporally structured communication at multiple scales. Our findings identify an infraslow modulation of hippocampal theta coherence that temporally organizes neuronal excitability across brain regions during REM sleep, offering a systems-level mechanism for regulating inter-regional communication during sleep.

## Introduction

Sleep is a fundamental brain state during which neuronal activity is reorganized across multiple temporal and spatial scales [1,2]. Classically, nREM sleep has been considered the canonical substrate for slow neural coordination, as it is dominated by slow oscillations, large-amplitude population synchrony, and prolonged periods of network stability [1,3,4]. These features make nREM sleep an ideal candidate for the emergence of slow modulatory processes that structure interregional communication over extended timescales. By contrast, REM sleep is characterized by activated cortical dynamics, sustained hippocampal theta oscillations, and elevated neuronal firing in the absence of overt behavior, features that have earned it the designation of paradoxical sleep [5–7]. Although REM sleep has long been implicated in memory processing, emotional regulation, and large-scale network integration [8,9], how hippocampal activity is temporally coordinated across distributed circuits during REM sleep remains incompletely understood.

The hippocampus is not a homogeneous structure, but instead exhibits pronounced functional differentiation along its septotemporal axis [10]. The dorsal hippocampus is preferentially engaged in spatial and cognitive processing, whereas the ventral hippocampus is more closely linked to affective and motivational functions [11]. Anatomically, these regions are interconnected yet receive non-overlapping inputs and project to different downstream targets, raising the possibility that their coordination is dynamically regulated rather than static [12]. Previous work has firmly established that the hippocampus is a theta oscillator, with theta oscillations propagating as a travelling wave, and exhibiting coherence along the septotemporal axis [6,13]. However, whether this coordination is stable or temporally structured, and whether such structure differs across sleep states, remains unknown.

Beyond fast oscillatory activity, growing evidence indicates that brain function is also shaped by very slow fluctuations on the order of tens of seconds to minutes. Indeed, infraslow rhythms (<0.1 Hz) have been observed in electrophysiological signals, neurovascular activity, and blood-oxygen-level–dependent signals, and have been linked to fluctuations in cortical excitability, behavioral performance, and functional connectivity [14–17]. These slow dynamics are increasingly recognized as an organizing scaffold that can modulate faster oscillations and neuronal firing. Given its prominent slow oscillatory activity, nREM sleep would be expected to provide a privileged context for such infraslow modulation. Nonetheless, the presence and functional relevance of infraslow dynamics within hippocampal circuits across distinct sleep states remain largely unexplored. Sleep provides a powerful framework for studying neural synchrony and large-scale coordination [1,2], as it encompasses discrete brain states with well-defined oscillatory regimes and minimal interference from ongoing behavior or sensory processing [18]. Across sleep stages, neuronal activity is reorganized over multiple temporal and spatial scales, offering a controlled context in which to examine how interregional communication emerges and is dynamically regulated. In particular, the relative stability of sleep states enables the expression of slow modulatory processes that are difficult to resolve during the rapid and variable transitions characteristic of wakefulness.

Within this framework, nREM sleep would be expected to dominate slow network modulation, whereas REM sleep has been primarily viewed through the lens of fast oscillatory coordination. However, REM sleep represents a uniquely activated yet behaviorally quiescent state, characterized by sustained hippocampal theta oscillations, elevated neuronal firing, and reduced sensory input. In addition, REM sleep engages widespread neuromodulatory systems and thalamo–cortical circuits that are well positioned to influence global excitability and synchrony across the forebrain [19–21]. Whether these features support slow, temporally structured modulation of hippocampal coordination has remained unclear.

In this study, we asked whether hippocampal coordination during sleep is dynamically modulated over extended timescales, and whether such modulation organizes neuronal activity across distributed hippocampal, cortical, and thalamic circuits. Using simultaneous recordings from such extended circuits in freely behaving rats, we examined oscillatory dynamics, interregional phase synchrony, and neuronal firing across the sleep–wake cycle. Contrary to the expectation that slow modulation would be most prominent during nREM sleep, we found that REM sleep uniquely amplifies theta coherence along the hippocampal septotemporal axis and reveals a pronounced infraslow modulation of this coupling. Importantly, this slow rhythm is associated with coordinated fluctuations in neuronal excitability across multiple brain regions, consistent with a network-level gain modulation mechanism. Together, these findings uncover an unexpected temporal structure of hippocampal coordination during REM sleep and provide a systems-level framework for understanding how REM sleep organizes large-scale communication across memory-related networks.

## Results

### Brain state-dependent hippocampal dynamics

To characterize hippocampal oscillatory dynamics across brains states, we performed simultaneous recordings of neuronal activity from anatomically interconnected hippocampal regions (CA1d and CA1v) and cortical areas (RSC and PFC) in rats across the sleep–wake cycle (Fig. S1, Table S1). In a subset of animals, recordings also included dorsal thalamic nuclei. During waking behavior, animals were allowed to explore a T-maze with return arms. Accordingly, two waking states were distinguished based on locomotion, running (RUN), when animals continuously traversed the maze, and quiet wakefulness (QW), defined by minimal movement (speed < 5 cm/s). For sleep recordings, animals were placed on a small platform and allowed to sleep ad libitum; and vigilance states were classified into non-REM (nREM) and rapid eye movement (REM) sleep using standard electrophysiological criteria.

Across states, hippocampal field potentials were dominated by two low-frequency rhythms; that is, delta (0.5–2 Hz) and theta (5–10 Hz) oscillations (**Fig. 1A–D**). During RUN, both CA1d and CA1v exhibited prominent theta-band activity, with theta power significantly higher in CA1d than CA1v (two-way ANOVA, P = 2.2x10^-58^, **Fig. 1E**). This activated state was accompanied by dense, temporally structured spiking across hippocampus, cortex, and thalamus, consistent with active sensorimotor engagement and decision-making demands. During QW, theta oscillations remained evident but were reduced relative to delta activity, while maintaining a power difference between hippocampal poles (**Fig. 1E**). Spiking activity persisted across regions but appeared less temporally organized, indicating a transition to an internally driven yet awake network state (**Fig. 1B**). During nREM sleep, hippocampal spectra were dominated by low-frequency activity with a broad delta peak and practically, abolition of theta power (**Fig. 1C**). In this state, CA1d and CA1v exhibited highly similar spectral profiles, consistent with strong global synchronization along the hippocampal septotemporal axis [22]. Correspondingly, neuronal firing across hippocampus, cortex, and thalamus was sparse and intermittent, reflecting downscaled network excitability [23,24]. In contrast, REM sleep was characterized by the re-emergence of a narrow, high-amplitude theta peak in both CA1d and CA1v, exceeding that observed during waking states and remaining consistently larger in CA1d (**Fig. 1E**). Despite the absence of overt behavior, REM sleep was associated with sustained spiking across hippocampal, cortical, and thalamic populations. Example activity traces illustrate continuous, highly regular theta oscillations, while spike rasters reveal dense yet state-specific firing patterns (**Fig. 1A-D**).

**Figure 1.**
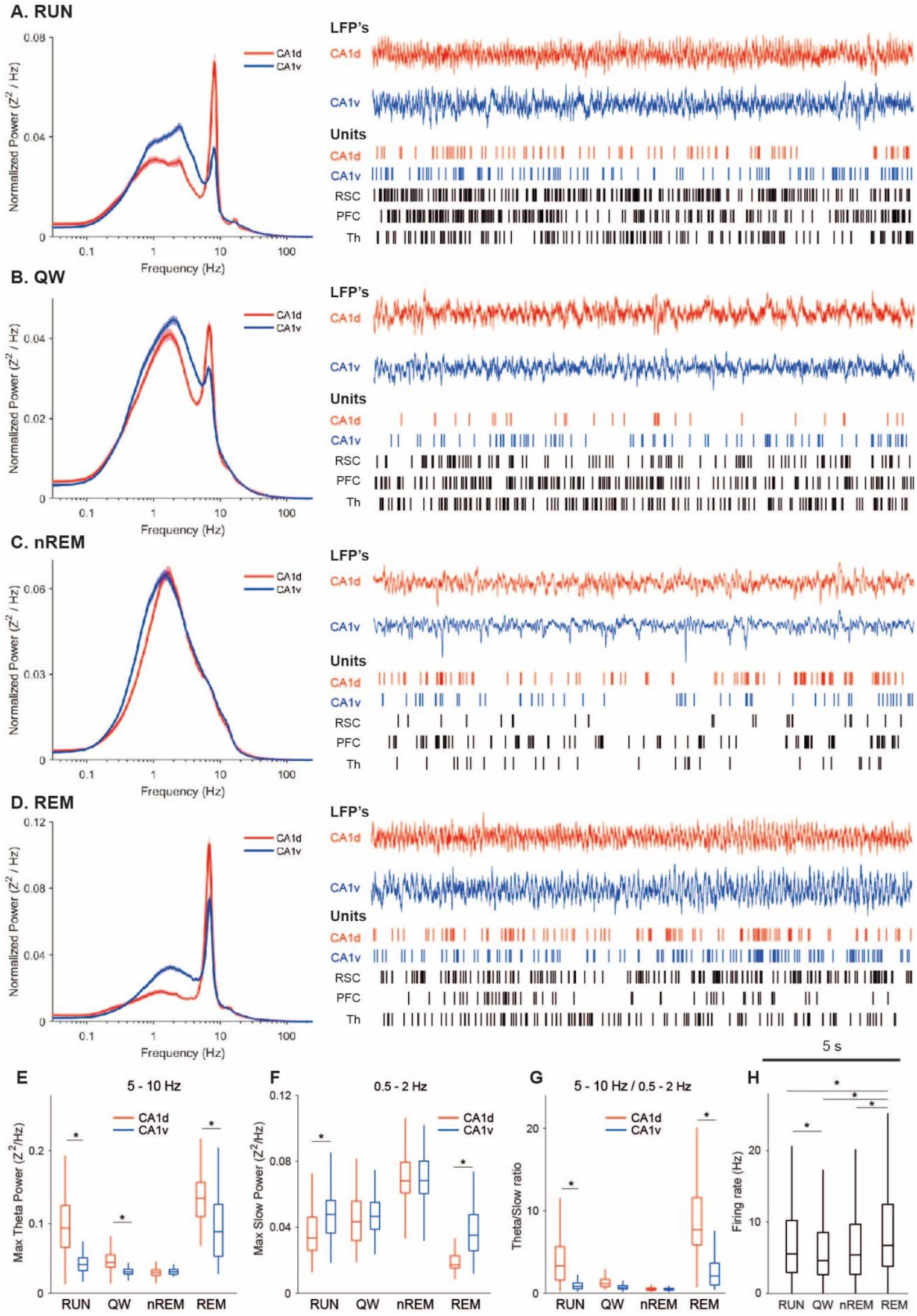
Brain state–dependent hippocampal oscillatory dynamics and distributed neuronal activity across the sleep–wake cycle. **A–D**, Examples of local field potentials (LFPs) and neuronal spiking recorded simultaneously from dorsal CA1 (CA1d), ventral CA1 (CA1v), retrosplenial cortex (RSC), prefrontal cortex (PFC), and dorsal thalamus (Th) during running (RUN), quiet wakefulness (QW), non–rapid eye movement sleep (nREM), and rapid eye movement sleep (REM). Left panels show normalized power spectral density of CA1d (red) and CA1v (blue) LFPs (logarithmic frequency axis). Right panels display example LFP traces (top) and spike rasters (bottom) from the same epochs. RUN and REM are characterized by prominent theta-band activity, whereas QW shows mixed theta–delta dynamics and nREM is dominated by low-frequency oscillations. During nREM sleep, neuronal firing across hippocampal, cortical, and thalamic regions is sparse and intermittent, whereas REM sleep exhibits sustained, state-specific spiking despite the absence of overt behavior. Scale bar, 5 s. **E**, Maximum theta-band power (5–10 Hz) in CA1d and CA1v across brain states. Theta power is highest during RUN and REM sleep and is consistently greater in CA1d than CA1v. **F**, Maximum slow/delta-band power (0.5–2 Hz) across states, peaking during nREM sleep and showing higher values in CA1v relative to CA1d. **G**, Theta-to-slow power ratio (5–10 Hz / 0.5–2 Hz) summarizing the net spectral shift across states. Activated states (RUN and REM) exhibit a pronounced increase in theta dominance, particularly in CA1d. **H**, Mean neuronal firing rate across recorded regions as a function of brain state. Firing rates are significantly elevated during REM sleep compared to RUN, QW, and nREM, indicating enhanced global excitability during this state. Box plots indicate median and interquartile range; whiskers denote data range. Asterisks indicate significant pairwise differences (^*^P < 0.05; see Tables S2–S6 for statistical details).

Quantification across animals confirmed that delta-band power peaked during nREM sleep (two-way ANOVA test, P = 7.4x10^-93^, Table S2), while theta-band power was maximal during REM sleep (two-way ANOVA, P = 2.6x10^-138^, Table S3). Delta power was consistently higher in CA1v than in CA1d (two-way ANOVA test, P = 1.3x10^-16^, **Fig. 1F**), whereas this pattern reversed in the theta band, with CA1d exhibiting stronger theta oscillations than CA1v (two-way ANOVA test, P = 2.2x10^-58^, **Fig. 1E**). To capture the net spectral shift across states, we computed the ratio of theta to delta power of CA1 regions for each state (**Fig. 1G**). This ratio was markedly elevated in CA1d relative to CA1v (two-way ANOVA test, P = 3.2x10^-36^), particularly during RUN and REM compared to QW and nREM (P < 0.001, Table S4), confirming that these activated states are defined by a shift toward theta-dominated dynamics. Finally, firing rate analysis revealed no interaction between hippocampal region and brain state (two-way ANOVA test, P = 0.055). Yet, CA1v consistently exhibited greater discharge than CA1d (P = 2x10^-7^), suggesting an excitability gradient along the septotemporal axis; and neuronal spiking was consistently higher during REM compared to the other brain states (Kruskal-Wallis, P =2.9x10^-5^, **Fig. 1H**, Table S5), pointing to an enhanced global neuronal excitability difference.

Together, these results confirm that hippocampal oscillatory structure and distributed neuronal discharge are strongly modulated by brain state. Theta oscillations dominate during activated states (RUN and REM sleep), particularly in CA1d, are attenuated during quiet wakefulness, and are suppressed during nREM sleep, with neuronal firing across hippocampal–cortical–thalamic circuits closely tracking these state-dependent oscillatory regimes.

### State-dependent theta coupling along the hippocampal septo-temporal axis reveals infraslow modulation during REM sleep

We next examined functional coupling along the hippocampal septotemporal axis by quantifying theta-band phase synchrony between CA1d and CA1v across brain states. Using the weighted phase-lag index (wPLI), we observed a prominent and narrowly tuned theta-band synchrony peak in all states (**Fig. 2A**). However, the magnitude of coupling was strongly modulated by brain state. Even during nREM sleep, when theta oscillations were largely suppressed, theta synchrony remained detectable, though at its lowest level. Theta coupling increased during QW and further during RUN, and reached its maximum during REM sleep, where wPLI values were consistently highest across animals (**Fig. 2A**).

**Figure 2.**
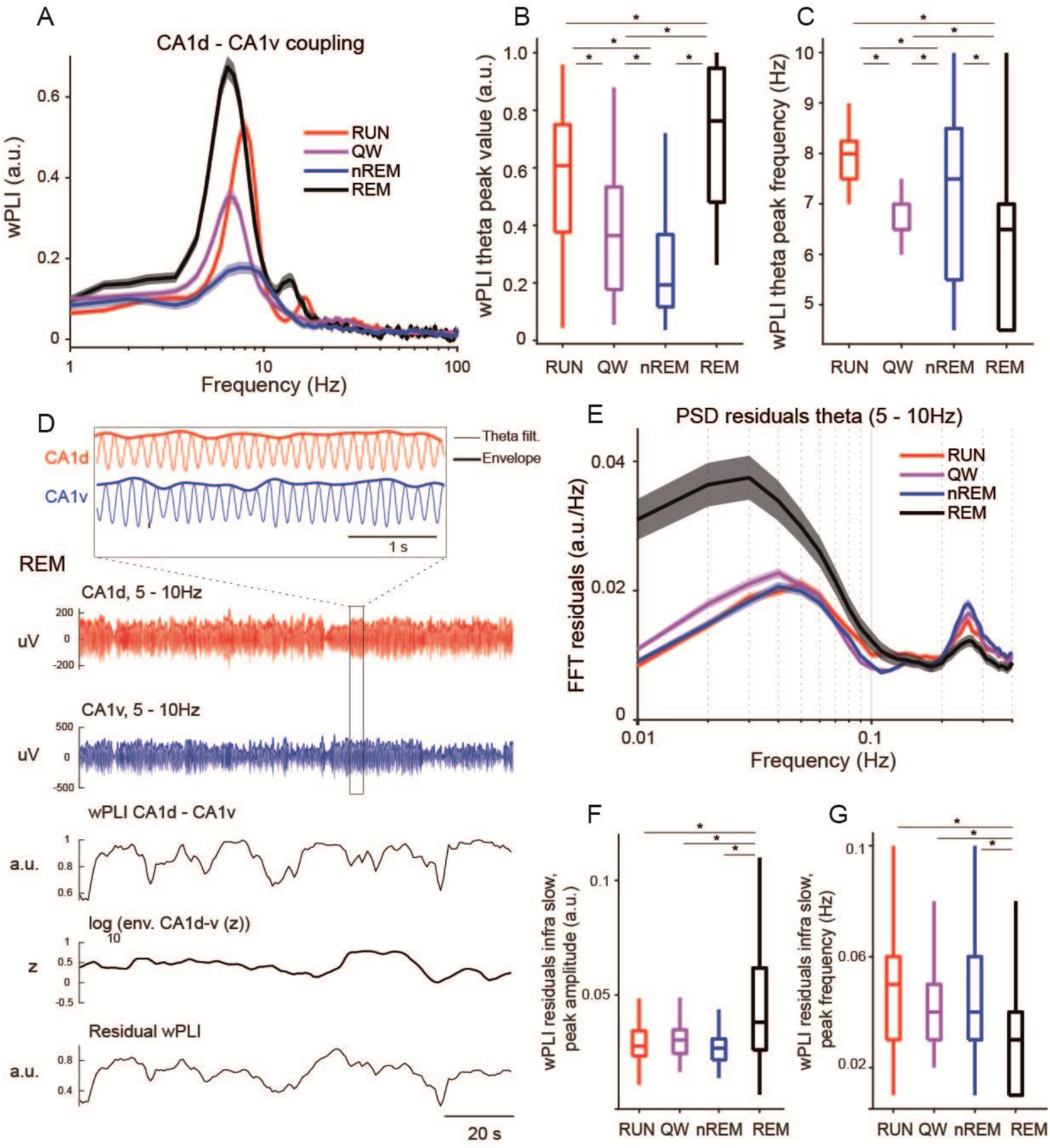
State-dependent theta coupling along the hippocampal septotemporal axis reveals infraslow modulation during REM sleep. **A**, Frequency-resolved weighted Phase Lag Index (wPLI) between CA1d and CA1v across brain states: running (RUN), quiet wakefulness (QW), non–rapid eye movement sleep (nREM), and rapid eye movement sleep (REM). All states exhibit a theta-band coupling peak, but coupling strength is markedly enhanced during REM sleep. Shaded areas indicate variability across animals. **B**, Peak theta-band wPLI value (5–10 Hz) across states. Theta coupling is strongest during REM sleep, intermediate during RUN and QW, and weakest during nREM sleep. **C**, Frequency at which theta-band coupling peaks across states. Peak coupling frequency varies modestly with brain state and is lowest during REM sleep. **D**, Example REM sleep epoch illustrating the temporal structure of theta coupling. Top inset shows band-pass filtered theta oscillations from CA1d and CA1v with their amplitude envelopes. Middle panels show raw theta-filtered LFPs (5–10 Hz) from CA1d and CA1v. Bottom panels show the time course of CA1d–CA1v wPLI, the log-transformed theta envelope (z-scored), and the residual wPLI after regression of theta power. The residual coupling signal exhibits slow fluctuations on the order of tens of seconds, independent of local theta amplitude. Scale bars as indicated. **E**, Power spectral density (PSD) of theta-band (5–10 Hz) power residuals across states. REM sleep exhibits a prominent infraslow peak (∼0.02– 0.03 Hz) that is strongly attenuated or absent in RUN, QW, and nREM sleep, indicating a REM-specific slow modulation of theta dynamics. **F**, Peak amplitude of the infraslow component in residual CA1d–CA1v wPLI. Infraslow modulation amplitude is significantly larger during REM sleep compared to all other states. **G**, Peak frequency of the infraslow modulation of residual wPLI across states. REM sleep is characterized by the slowest modulation frequency, consistent with a state-dependent gain modulation rather than a shift in theta frequency. Box plots indicate median and interquartile range; whiskers denote data range. Asterisks indicate significant pairwise differences (^*^P < 0.05).

Quantification of the theta-band peak synchrony confirmed a robust state dependence, with REM sleep exhibiting the strongest synchrony, followed by RUN, then QW, and nREM showing the weakest synchrony (**Fig. 2B**). The frequency at which theta coupling peaked varied modestly across states but remained within a narrow range (approximately 5–9 Hz) and it was lowest during REM (**Fig. 2C**). Importantly, linear mixed-effects modeling indicated that peak frequency did not predict coupling strength, suggesting that state-dependent differences primarily reflect modulation of interregional phase coordination (type III ANOVA marginal test, P = 7.7x10^-79^) rather than systematic shifts in theta frequency (type III ANOVA marginal test, P = 0.35).

Inspection of the temporal evolution of CA1d–CA1v theta synchrony revealed that coupling strength was not stationary but fluctuated rhythmically over time (**Fig. 2D**). These fluctuations were particularly pronounced during REM sleep, where the wPLI time series exhibited clear infraslow oscillations on the order of tens of seconds. Crucially, this structure persisted after regressing out theta-band power, indicating that the observed infraslow modulation reflects genuine changes in interregional phase coupling rather than trivial power covariation (Fig. S2, **Fig. 2D**). Similar infraslow fluctuations were observed in the theta-band amplitude envelope, pointing to a shared modulatory process influencing both local oscillatory power and dorso-ventral hippocampal synchrony.

To further characterize the infraslow modulation, we computed the spectral content of theta-band power residuals after removing state-specific mean power profiles (**Fig. 2E**), notwithstanding that the correlation between raw wPLI and theta power was generally low across the population (median *R*^*2*^ ∼ 0.1; see Fig. S3). Across animals, REM sleep exhibited a pronounced infraslow peak centered near 0.02–0.03 Hz, which was largely absent or markedly reduced during all other brain states. This infraslow component was likewise present in residual CA1d–CA1v wPLI time series, where REM sleep showed a significantly larger infraslow peak amplitude than all other states (**Fig. 2F**). Moreover, the peak frequency of this infraslow modulation was slowest during REM sleep (**Fig. 2G**), consistent with a state-dependent gain modulation rather than a simple frequency shift. Furthermore, we examined whether comparable infraslow dynamics were also present in cortical connectivity. To this end, we computed the spectral content of the residual theta-band power for RSC–PFC interactions (Fig. S3). The peak infraslow modulation observed in the cortex was significantly smaller than that detected in the hippocampus (Wilcoxon rank-sum test, P = 1.68x10^-8^), indicating that this infraslow dynamic modulation predominantly affected hippocampal activity rather than cortical networks.

Together, these results demonstrate that theta-band coupling along the hippocampal septotemporal axis is dynamically regulated by brain state. While theta synchrony is present during all brain states, REM sleep uniquely amplifies both the strength and temporal variability of CA1d–CA1v coupling, introducing a prominent infraslow modulation that coordinates local theta power and interregional phase coupling. This finding extends the state-dependent spectral reconfiguration described above by revealing an additional, slower temporal structure that selectively governs hippocampal communication, preferentially during REM sleep.

### Temporal evolution of infraslow hippocampal coordination during brain state transitions

We next investigated how hippocampal theta coupling and multi-regional spiking activity evolve around transitions between brain states, focusing on the infraslow modulation observed during REM sleep. We first aligned CA1d–CA1v theta synchrony and multi-unit activity (MUA) across hippocampal, cortical, and thalamic regions to the onset of REM sleep (**Fig. 3A**). This analysis revealed a marked increase in hippocampal theta coupling that began shortly before REM onset and persisted throughout the REM episode. Concurrently, firing rates gradually rose across all recorded regions, with the thalamus exhibiting the most prominent activation. These coordinated changes suggest that REM onset is characterized by a synchronous increase in both interregional theta phase alignment and global neuronal excitability. To quantify this effect, we examined wPLI trajectories across transitions into REM, nREM, and QW. Theta coupling during REM showed a sustained increase over time, whereas transitions into nREM or QW were associated with declining synchrony (**Fig. 3B**). Estimating the initial slope of the wPLI signal following each transition confirmed that only REM onset produced a significant positive ramp in theta coupling (**Fig. 3C**), reinforcing the notion that REM initiates a structured gain modulation of theta coherence across the septotemporal axis of the hippocampus.

**Figure 3.**
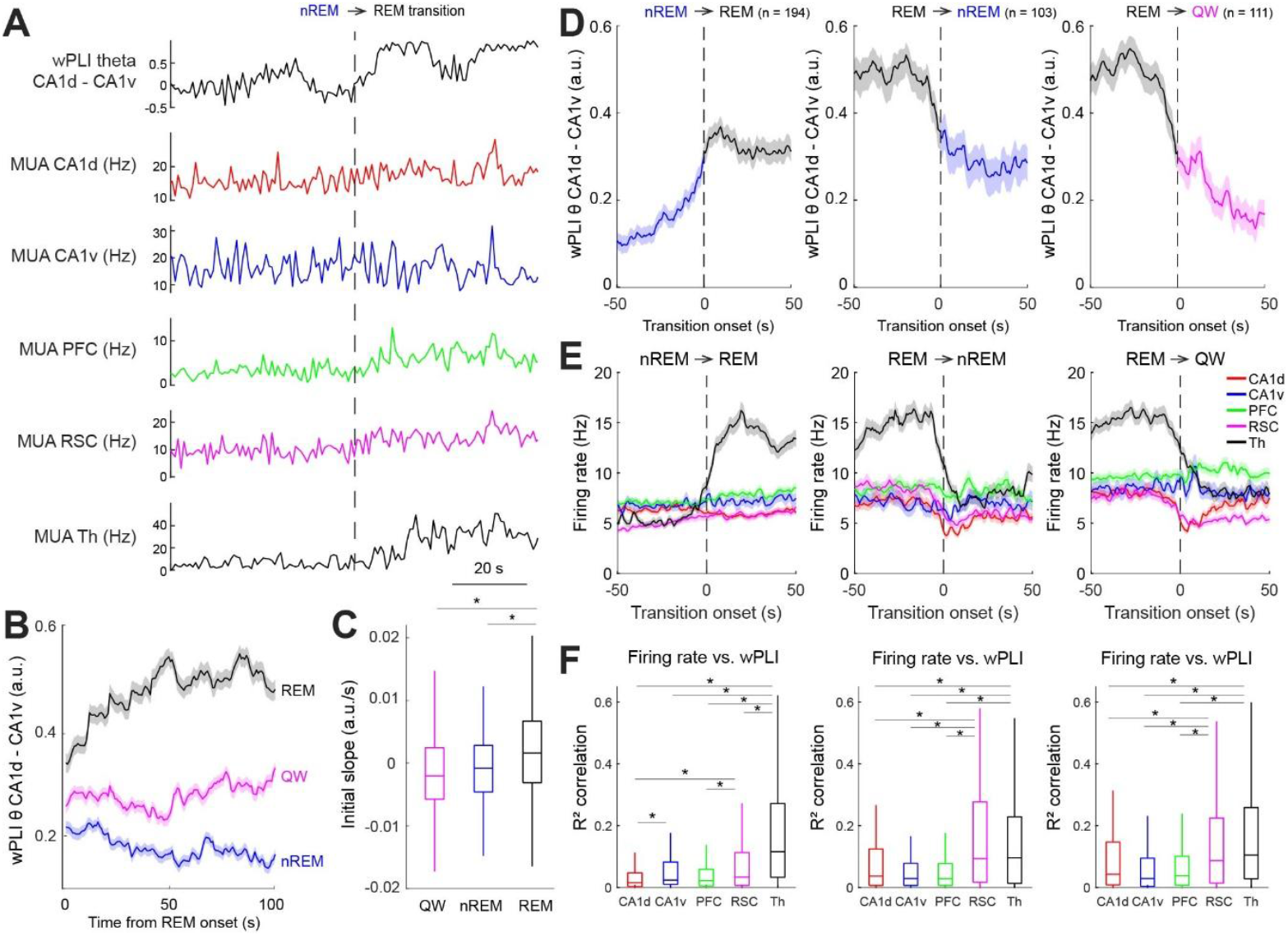
Temporal evolution of hippocampal theta coupling and distributed spiking activity during brain state transitions. **A**, Example transition from nREM to REM sleep illustrating the time course of CA1d–CA1v theta-band coupling (wPLI; top) and multiunit activity (MUA) across dorsal CA1 (CA1d), ventral CA1 (CA1v), prefrontal cortex (PFC), retrosplenial cortex (RSC), and dorsal thalamus (Th). Signals are aligned to REM onset (dashed line). Theta coupling and firing rates increase gradually around the transition, with the strongest activation observed in the thalamus. **B**, Mean wPLI trajectories aligned to REM onset for transitions into REM (black), quiet wakefulness (QW; magenta), and nREM sleep (blue). Theta coupling increases selectively during REM episodes and remains comparatively low during QW and nREM. **C**, Initial slope of the wPLI signal following transition onset. Only transitions into REM sleep exhibit a significant positive slope, indicating a gradual buildup of hippocampal theta synchrony specific to REM onset. **D**, Average CA1d–CA1v theta coupling aligned to different transition types: nREM → REM, REM → nREM, and REM → QW. Coupling increases during nREM → REM transitions and decreases sharply at REM offset, regardless of whether the subsequent state is nREM or QW, revealing an asymmetric regulation of hippocampal synchrony across REM boundaries. Shaded areas indicate SEM; numbers denote transition counts. **E**, Mean firing rate dynamics across regions aligned to the same transitions shown in (D). Thalamic firing exhibits the most pronounced increase during nREM → REM transitions and the sharpest decrease at REM termination, whereas hippocampal and cortical regions show more moderate changes. **F**, Coefficient of determination (R^2^) between regional firing rates and CA1d–CA1v theta coupling across time for each transition type. Thalamic activity shows the strongest correlation with coupling dynamics, indicating that thalamic excitability best predicts fluctuations in hippocampal theta synchrony. Box plots show median and interquartile range; whiskers denote data range. Asterisks indicate significant differences (^*^P < 0.05).

Further parsing transitions specifically involving REM (nREM→REM, REM→nREM, REM→QW) revealed a clear asymmetry, so that theta coupling reliably increased during nREM→REM transitions, but decreased during REM offsets, regardless of whether the animal transitioned into nREM or QW (**Fig. 3D**). This pattern underscores that REM sleep is associated with a state-dependent upregulation of hippocampal theta synchrony that is rapidly reversed as the network exits REM sleep. These dynamics indicate that CA1d–CA1v coordination is under tight temporal control, with REM sleep uniquely driving a reversible elevation in septotemporal theta coherence.

To further understand how these changes relate to regional firing dynamics, we aligned MUA traces to the same transition events (**Fig. 3E**). During nREM→REM transitions, firing rates increased dramatically in the thalamus. Upon REM termination, thalamic activity sharply declined, while other regions exhibited more modest reductions. Indeed, CA1v and PFC activity remained relatively stable across transitions, whereas CA1d and RSC showed modest but detectable declines. This divergence suggests that the thalamus plays a pivotal role in orchestrating global state transitions and may act as a central driver of REM-associated network reconfiguration. To probe this relationship, we computed the coefficient of determination (R^2^) between regional firing rates and CA1d–CA1v theta coupling across time (**Fig. 3F**). Thalamic activity exhibited the highest correlation with theta coupling dynamics, suggesting that this region is particularly influential in modulating hippocampal synchrony. In contrast, hippocampal and prefronta firing rates were weakly associated with coupling fluctuations. These findings imply that the slow buildup of theta synchrony during REM sleep is more closely linked to changes in thalamo-cortical excitability than to local hippocampal firing.

In sum, these results demonstrate that REM sleep is characterized by a distinct temporal regime in which hippocampal theta coordination and distributed spiking activity coordinately evolve over infraslow timescales. REM onset triggers a gradual but structured increase in theta synchrony along the hippocampal septotemporal axis, accompanied by widespread elevations in neuronal excitability, especially in the thalamus. This configuration is reversed at REM offset, highlighting the transient and state-specific nature of the REM network. The strong correlation between thalamic spiking and hippocampal coupling points to a hierarchical mechanism in which REM-related gain modulation of theta synchrony is driven by upstream excitability changes.

### Network phase modulation of neuronal spiking by infraslow hippocampal synchrony

To investigate whether the infraslow modulation of CA1d–CA1v theta coupling during REM sleep reflects structured fluctuations in neuronal excitability, we examined its relationship with spiking activity across hippocampal, cortical, and thalamic regions. We first pooled all units across regions and aligned their activity to the phase of the infraslow theta synchrony signal, separately for each brain state. This analysis revealed distinct phase preferences across states, suggesting that the timing of neuronal excitability relative to hippocampal theta coordination is state-dependent (Fig. S4). Next, we aligned the wPLI time series to simultaneously recorded MUA during REM sleep (**Fig. 4A**). Slow oscillations in theta coupling on the order of tens of seconds were clearly visible, and these fluctuations coincided with rhythmic modulations of spiking across the circuit. Notably, transient peaks in CA1d–CA1v theta synchrony were accompanied by elevated MUA in all recorded regions, suggesting that theta coupling dynamics reflect a shared excitability signal that temporally coordinates distributed neuronal activity. To determine whether regional firing was phase-locked to the infraslow theta synchrony signal, we extracted the instantaneous phase of the wPLI trace and computed phase locking distributions. Circular histograms and Rayleigh tests confirmed significant phase locking in all regions (non-uniformity Rayleigh test, p < 0.001, **Fig. S4**), with concentrated unimodal distributions and strong resultant vectors. Although preferred phases varied slightly between areas, they generally clustered near the trough of the theta coupling cycle, consistent with a common infraslow modulation across the hippocampo–cortical– thalamic loop.

**Figure 4.**
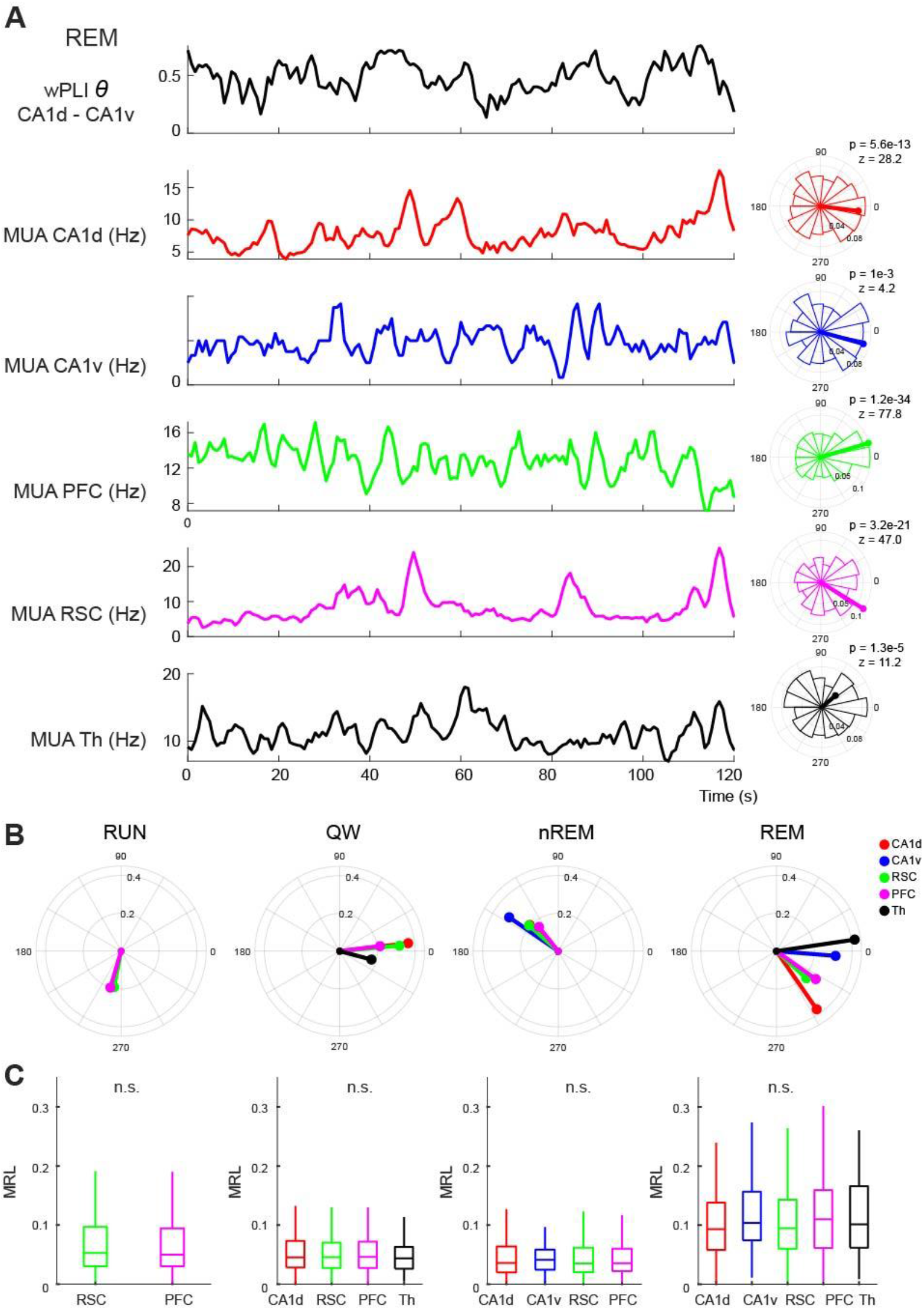
Infraslow modulation of hippocampal theta coupling entrains neuronal firing across hippocampal–cortical–thalamic circuits during REM sleep. **A**, Example REM sleep episode illustrating the temporal relationship between infraslow fluctuations in CA1d–CA1v theta coupling (wPLI; top trace) and multiunit activity (MUA) recorded from dorsal CA1 (CA1d), ventral CA1 (CA1v), prefrontal cortex (PFC), retrosplenial cortex (RSC), and dorsal thalamus (Th). Traces are aligned in time (bottom axis). Right panels show circular histograms of spike phases relative to the instantaneous phase of the infraslow theta coupling signal for each region. Rayleigh test statistics (Z) and corresponding p values are indicated, demonstrating significant phase locking of neuronal firing to the infraslow modulation during REM sleep. **B**, Mean phase vectors for each recorded region during running (RUN), quiet wakefulness (QW), non–rapid eye movement sleep (nREM), and REM sleep. Vector direction indicates preferred phase of firing relative to the infraslow theta coupling cycle, and vector length reflects the strength of phase locking (mean resultant length, MRL). Consistent and directionally aligned phase preferences across regions emerge selectively during REM sleep. **C**, Population summary of mean resultant length (MRL) across regions and brain states. Phase locking is weak and not significantly different across regions during RUN, QW, and nREM, whereas REM sleep shows uniformly elevated MRL values across hippocampal, cortical, and thalamic regions, indicating robust and distributed entrainment to the infraslow modulation of hippocampal theta coupling. Box plots show median and interquartile range; whiskers denote data range. “n.s.” denotes non-significant differences across regions.

We next assessed whether such phase locking was specific to REM sleep by computing mean phase vectors for each region across brain states (**Fig. 4C**). During RUN, only cortical units showed significant modulation, with no difference in phase alignment between RSC and PFC (Watson-Williams multi-sample test, P = 0.096). In QW, all regions except CA1v exhibited phase locking near a common phase (Watson-Williams multi-sample test, P = 0.15). During nREM sleep, thalamic units were not significantly modulated, whereas the remaining regions showed overlapping phase preferences (Watson-Williams multi-sample test, P = 0.48). In contrast, REM sleep was marked by robust and directionally aligned phase vectors in all regions (Watson-Williams multi-sample test, P = 1.25x10^-11^, Table S6), indicating network entrainment to the infraslow theta coupling signal. To quantify the consistency of this modulation, we calculated the mean resultant length (MRL) for each region and compared values across states (**Fig. 4D**). MRL values remained low during AE, QW, and nREM, indicating weak or inconsistent modulation. In REM sleep, MRL increased across all areas, reflecting stronger entrainment, although variability across animals limited statistical significance at the group level. These results suggest that while the strength of modulation may fluctuate across sessions, the infraslow rhythm reliably structures firing within individual REM episodes.

In summary, these findings demonstrate that the infraslow modulation of hippocampal theta coupling during REM sleep is closely linked to network fluctuations in excitability. This rhythm synchronizes neuronal firing across hippocampal, cortical, and thalamic structures in a temporally organized manner, supporting the notion that REM sleep engages a slow, gain modulation mechanism operating at the level of the entire network. By coordinating neuronal activity at multiple timescales, this infraslow dynamic may facilitate selective routing of information and temporally gate hippocampal–cortical communication.

## Discussion

Our study provides converging evidence that REM sleep orchestrates a unique mode of hippocampal and distributed thalamo-cortical coordination, characterized by theta-dominated oscillatory dynamics, heightened spiking activity, and infraslow modulation of interregional coupling. By leveraging simultaneous recordings from hippocampal, cortical, and thalamic circuits, we uncovered a temporally layered architecture of connectivity changes across brain states, highlighting REM sleep as a state ant the hippocampus as a hub of heightened infraslow synchrony.

REM sleep has long been associated with elevated hippocampal theta oscillations [6,25,26], but the circuit-level organization and temporal modulation of these dynamics have remained incompletely characterized. Our findings confirm and extend this classical view by demonstrating that theta oscillations are not only strongest during REM, but that CA1d and CA1v show increased coherence specifically during this state [22,27,28]. Furthermore, the emergence of an infraslow modulation (∼0.02 Hz) of theta synchrony during REM sleep reveals an additional regulatory layer that structures circuit excitability over extended timescales. This rhythm was largely absent in other vigilance states and cortical regions, indicating that REM sleep induces a temporally unique coordination mode. Importantly, the presence of REM-specific infraslow fluctuations was detected in the dorso-ventral hippocampus phase synchrony, not in the local field potential power, suggesting the engagement of a global modulatory mechanism [14,29]. While this signal may arise from neurovascular rhythms, such as vasomotion [30,31], that exhibit similar temporal structure and widespread spatial reach, our data indicate that its influence extends into neuronal spiking. Indeed, neurons across hippocampal, cortical, and thalamic regions showed significant phase locking to the infraslow rhythm derived from theta coherence, indicating that the infraslow fluctuation is functionally relevant and capable of shaping circuit excitability [32,33]. Notably, this modulation was most consistent during REM sleep, supporting the idea that infraslow coupling serves as a gain-modulating scaffold for hippocampo–cortical–thalamic interactions, particularly during this state.

These results align with and extend prior work in humans and rodents showing that REM sleep engages unique oscillatory modes [26,34]. Human EEG and intracranial recordings have identified transient increases in theta and gamma power during REM [26], with proposed roles in emotional memory processing and sensorimotor integration. Our data add a systems-level perspective by showing that these faster rhythms are embedded within a slower temporal scaffold that modulates their expression and interregional coordination. This temporal nesting may provide a mechanism by which REM sleep supports both local plasticity and distributed integration across memory networks [35,36]. The tight coupling between thalamic spiking and theta synchrony further points to a key role for thalamo–cortical circuits in driving or coordinating REM-related network states. Transitions into REM sleep were marked by pronounced increases in thalamic firing, which most significantly predicted increases in CA1d–CA1v theta coherence. This suggests that thalamic excitability may gate hippocampal communication or facilitate large-scale synchronization across the forebrain [1,37]. These findings echo earlier work showing that the midline thalamus is essential for REM sleep generation and supports memory-dependent communication between the hippocampus and PFC [37,38].

Interestingly, we also observed state-specific dissociations in spiking activity across hippocampal fields. CA1v exhibited higher firing rates than CA1d across all states, yet theta power and coherence were consistently stronger in CA1d. This suggests that theta rhythmicity and firing rate are dissociable features, possibly reflecting distinct roles of dorsal and ventral hippocampus in REM-related computations. Previous work has shown that CA1d is more involved in spatial encoding, while CA1v contributes to affective processing and motivational drive [10]. The differential coordination of these fields during REM could reflect their engagement in distinct, yet complementary, consolidation processes. Our results also highlight the temporal asymmetries of REM transitions. The onset of REM was associated with a gradual increase in theta coherence and spiking activity, while transitions out of REM (into either nREM or QW) exhibited sharp decreases. This asymmetry suggests that REM onset may involve a progressive build-up of synchrony and excitability, potentially facilitating memory reactivation or integration. By contrast, REM offset may represent a rapid disengagement from this integrative mode, consistent with the functional segmentation of sleep stages [18,39].

In addition to REM-specific dynamics, our data also revealed important contrasts across other states. During RUN, theta oscillations were prominent but lacked the infraslow modulation seen in REM, indicating that active wakefulness supports strong but more stationary interregional coordination. During QW and nREM, theta power and coherence declined, and spiking became sparser and less structured. These observations confirm that hippocampo–cortical–thalamic communication is tightly state-gated [1] and that REM occupies a privileged position in the oscillatory hierarchy. One important implication of these findings is that REM sleep may support temporally segmented memory processing through infraslow modulation. Rather than maintaining constant high synchrony, the theta coherence signal oscillates on a 30–50 s timescale, creating alternating windows of high and low excitability. These windows could gate the reactivation of memory traces or the integration of mnemonic content across regions. Similar mechanisms have been proposed in human studies linking infraslow rhythms to fluctuations in cognitive performance and BOLD signals during rest [14,15,17]. Despite these insights, several limitations should be acknowledged in our study. First, the sample size was moderate, and although consistency across animals was high, future work with larger cohorts will be necessary to confirm the generalizability of these findings. Second, we did not directly test memory retention following REM episodes with specific theta coherence profiles. While the association between REM and certain types of memory consolidation is well established, demonstrating causal links between the infraslow modulation described here and behavioral outcomes will require targeted perturbation experiments. Third, the use of anesthetized or head-fixed animals in some studies of REM coordination may differ from our freely behaving setup; thus, comparisons should be interpreted cautiously. Fourth, we cannot fully rule out the presence of the infraslow oscillation during active behavior (RUN). The primary limitation lies in its extended timescale (30–50 s), which exceeds the duration over which animals typically sustain a consistent behavioral state during exploration. As a result, detecting slow fluctuations is confounded by frequent behavioral transitions. Future experimental designs with more stable behavioral epochs will be necessary to determine whether this infraslow modulation also emerges during active waking states.

Moreover, future work should explore whether the infraslow modulation of hippocampal coupling is altered in models of cognitive dysfunction or neuropsychiatric disease. Disruptions of REM sleep and theta rhythms have been implicated in conditions such as depression, schizophrenia, and Alzheimer’s disease [40–42]. If the infraslow modulation we describe is critical for coordinating distributed memory networks, its breakdown may contribute to the cognitive impairments observed in these disorders. Additionally, studies in humans using MEG or high-density EEG may be able to identify similar rhythms, providing translational bridges across species [43]. Mechanistic studies are needed to determine the origin of the infraslow signal. While vasomotor activity is a plausible contributor, the observed coupling to spiking suggests that neuromodulatory inputs, such as cholinergic or noradrenergic tone, may also play a role [20,44]. Simultaneous recordings of vascular signals, field potentials, and neuromodulator release could disentangle these contributions [31].

In summary, our results reveal that REM sleep induces a distinct hippocampal coordination mode characterized by high theta power, elevated coherence between CA1 fields, and a REM-specific infraslow modulation that structures excitability across the hippocampo–cortical–thalamic network. This layered dynamic scaffolding may support temporally segmented consolidation processes over prolonged cycles and provides a novel systems-level framework for understanding REM sleep’s role in memory and brain-wide communication.

## Methods

### Animals

Fifteen adult male Sprague–Dawley rats (postnatal days 40–60; 300– 350 g) were obtained from the Center for Innovation on Biomedical Experimental Models (CIBEM, Pontificia Universidad Católica de Chile). Animals were housed under controlled temperature (22 ± 1 °C) and a 12 h light/dark cycle (lights on at 08:00), with food and water available ad libitum. All experimental protocols were reviewed and approved by the Scientific Ethical Committee for the Care of Animals and the Environment of the Pontificia Universidad Católica (CEC-CAA), under approval number, protocol 220512003. All methods were carried out in accordance with relevant guidelines and regulations. This study is reported in compliance with the ARRIVE (Animal Research: Reporting of In Vivo Experiments) guidelines. Two weeks before surgery, animal handling was performed for 10 minutes daily for 3-5 consecutive days.

### Habituation

Animals were habituated to the housing environment for three days, then handled daily by the experimenter for three to five additional days. Subsequently, rats were habituated to the experimental environment by placing them on a towel-covered drum in the recording room for 30–60 minutes daily over approximately one week. Habituation to the T-maze with return arms was conducted over 3–5 consecutive days, with daily sessions ranging from 10 to 30 minutes to ensure adequate exploration and reduction of novelty-induced stress.

### Recording Implant Assembly

The recording drive was designed using Autodesk Fusion software and 3D-printed on a Fusion360 F410 printer. Sixteen or 32 tungsten standard tapered-tip electrodes (MicroProbes, USA; impedances of 1, 3, and 5 MΩ) were assembled onto a custom-fabricated drive targeted to brain regions of interest based on a rat brain stereotaxic atlas[45]. Each electrode was connected to an EIB-18 or EIB-36 PCB card (capacity of 18 or 36 channels, including ground and reference). Ground and reference channels were connected to stainless steel screws placed into the skull during surgery. The entire assembly was covered with copper mesh to minimize electrical noise.

### Stereotaxic Surgery

Rats were anesthetized with isoflurane (4% induction, 1.5–2% maintenance) and secured in a stereotaxic frame (Stoelting Inc.). Body temperature was maintained at 35–37 °C using a homeothermic blanket, and animals received hourly hydration with glucosaline solution (0.9% NaCl, 2.5% dextrose). Following a scalp incision, four craniotomies (∼1 mm diameter each) were performed in the right hemisphere at predetermined stereotaxic coordinates (CA1d: -4.0 AP, 2.0 ML, 2.7 DV; CA1v: -5.7 AP, -5.5 ML, 7.2 DV; RSC: -4.0 AP, 0.7 ML, 2.2 DV; PFC: 2.5 AP, -0.7 ML, 4.2 DV; Th: -2.1 AP, 0.7 ML, 5.2 DV, all in mm). Two additional craniotomies anterior to bregma accommodated the ground and reference electrodes, while four more craniotomies (two parietal contralateral, one parietal ipsilateral, and one posterior to lambda) secured the implant with additional screws. The dura was carefully removed in the craniotomies, cortical surfaces were moistened with mineral oil, and electrodes were carefully inserted. Craniotomies were sealed with silicone elastomer or wax, and the recording drive was fixed using dental acrylic. Postoperative care involved daily subcutaneous injections of enrofloxacin (10 mg/kg) and meloxicam (1 mg/kg) for three consecutive days. Animals recovered for at least seven days before recordings began.

### Electrophysiological Recordings

Following recovery, electrophysiological recordings were conducted over 10 consecutive days during the light phase in a Faraday-shielded enclosure. Rats performed a self-paced two-armed bandit in a maze with a start box, a central stem, and two lateral goal ports (nose-pokes). Periods of active locomotion, defined by a running speed exceeding 5 cm/s, were identified as RUN epochs and extracted for analysis. Subsequently, rats were placed on a towel-covered platform for 60–90-minute sessions. The EIB-18 or EIB-36 board was connected via a 16- or 32 channel headstage (Intan Technologies) to an amplifier (Intan RHD Recording System, Intan Technologies, USA), with one electrode serving as reference. Video recordings synchronized with the amplifier clock aided in brain state identification. Signals were sampled at 20 kHz using RHX Data Acquisition software (Intan Technologies) and later converted to MATLAB format using the LAN toolbox for further analysis.

### Histology

After the recording protocol finished, rats were anesthetized with isoflurane (4% induction, 1.5–2% maintenance), and electrolytic lesions (5 µA for 10 s) marked electrode locations. After 48 h of recovery, animals were anesthetized to a surgical plane with ketamine (300 mg/kg) and xylazine (30 mg/kg, intraperitoneally [i.p.]) and euthanized by transcardial perfusion with 0.9% saline followed by 4% paraformaldehyde. Brains were postfixed overnight, transferred to PBS-azide, and sectioned coronally using a vibratome (World Precision Instruments, USA). Sections were Nissl-stained and examined with a Nikon Eclipse CI-L microscope to verify electrode placements.

### Sleep scoring

CA1d LFP was downsampled and time-frequency decomposition was performed with Fourier analysis using the LAN toolbox to obtain the signal’s power spectral density. The signal was analyzed for subsequent 10-s windows, where the raw LFP signal, the power spectrum, and the video recording were used to determine the stage of the sleep-wake cycle. One of three different stages was assigned to each window depending on the following criteria: if the power spectrum had a peak in slow wave activity (0.5–4 Hz) and the animal was immobile, the window was identified as NREM; if the power spectrum presented a peak in theta oscillations (4–10 Hz) and the animal was immobile, the window was identified as REM sleep; finally, if the animal was actively moving, the window was identified as quiet wakefulness (QW). To assign a window into any of these categories, the criteria had to be fulfilled in at least 50% of the window. Exceptionally, a window was scored as undetermined if the previously described criteria were not met. This analysis was performed with a MATLAB (MathWorks, Natick, MA) script.

### Spike Sorting

Extracellular signals were band-pass (600-5000 Hz) filtered and the spike waveforms with either negative or positive peaks exceeding were extracted. Single units were sorted using a custom manual clustering program (Kilosort2, https://github.com/MouseLand/Kilosort) and were distinguished based on principal components. Putative pyramidal cells and interneurons were further separated by principal component analysis of the unit waveforms and mean firing rat. A unit cluster was classified as a single-unit activity if the refractory period (time between two consecutive spikes) was at least 1.5 ms. When the recording quality and the spike sorting did not allow unambiguous single-unit isolation, the spike cluster was conservatively classified as multiunit activity (MUA).

### Spectral analysis

Time-frequency decomposition of LFP was performed with multi-taper method implemented in the Chronux toolbox (http://www.chronux.org, function *mtspecgramc*) for MATLAB. Analysis was performed on signals downsampled to 500 Hz. We used a sliding window of 30 s with a 1 s step size. The multi-taper parameters were configured with a time-bandwidth product of 5 and 9 tapers, providing an optimized balance between spectral resolution and variance reduction. The frequency range of interest was restricted to 0-245 Hz (zero padding = 0). The representative PSD profile was obtained by calculating the median power across all time windows to minimize the impact of transient artifacts and non-stationary noise.

Complementary, after calculating the weighted Phase Lag Index (wPLI) between signal pairs with a temporal resolution of 0.8 s we evaluated the frequency components of the wPLI residuals for theta band (see *Weighted phase lag index analysis* for details) using Welch’s power spectral density estimate (*pwelch* function in MATLAB) to identify periodicities within the connectivity fluctuations.

### Weighted phase lag index (wPLI) analysis

To evaluate the functional coupling between LFP signals, we calculated the debiased Weighted Phase Lag Index (wPLI) using the FieldTrip toolbox (https://www.fieldtriptoolbox.org). This metric was selected for its robustness against volume conduction and its reduced sensitivity to sample size bias. The continuous LFP data were analyzed using a sliding window approach with a 10 s window length and a 0.8 s step size. Within each 10 s window, signals were z-score normalized and further segmented into 2 s sub-epochs. For each sub-epoch, the cross-spectral density was estimated using a fast Fourier transform (FFT) with a multitaper approach (Discrete Prolate Spheroidal Sequences, DPSS). The frequency range was restricted to 1 – 100 Hz. The wPLI debiased estimator was then computed across the sub-epochs to obtain a single connectivity value per time step. This procedure yielded a time-resolved wPLI spectrum, representing the phase-lead/lag relationship between the two recording sites while effectively minimizing the contribution of zero-lag noise.

To ensure that the observed fluctuations in wPLI were not driven by changes in signal amplitude (the “power effect”), we performed a residualization analysis using a multiple linear regression model. First, we extracted the mean wPLI values within the theta band (5–10 Hz). Simultaneously, we calculated the instantaneous power envelope for both LFP channels (LFP1 and LFP2) by applying a 4th-order Butterworth bandpass filter followed by a Hilbert transform. The resulting envelopes were log-transformed, z-score normalized, and smoothed using a moving average filter to match the temporal scale of the connectivity windows. These power traces were then resampled to the wPLI time axis via linear interpolation. Finally, we implemented a multiple linear regression where the theta-band wPLI was the dependent variable, and the normalized power envelopes of both LFP1 and LFP2 served as predictors. The residual signal (*g*) was defined as the component of the wPLI that could not be explained by the linear combination of the local power fluctuations:

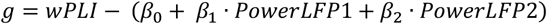

where, *β*_1_ and *β*_2_ represent the regression coefficients that quantify the specific contribution of the power fluctuations of LFP1 and LFP2, respectively, to the variance of the wPLI signal. By subtracting these terms, the residual wPLI (*g*) provides a measure of coupling dynamics that is independent of local power changes. This residual signal was used for subsequent frequency decomposition to identify intrinsic periodicities in the synchronization between both regions.

### Spike-phase coupling

Theta band wPLI time series was first band-pass filtered within infra-slow band using a high-order Butterworth filter implemented in a zero-phase forward-backward scheme (*filtfilt* function in MATLAB) to avoid temporal shifts. The instantaneous phase of the filtered wPLI was then extracted using the Hilbert transform. Specifically, the analytic signal was computed to obtain the unwrapped phase trajectory. For each recorded neuron, we extracted the corresponding wPLI phase at the exact timestamps of its spikes.

### Statistical Analysis

Group differences with a single, independent factor were examined with a one-way ANOVA when the data met parametric assumptions (normality and homogeneity of variances); if normality was violated the non-parametric Kruskal–Wallis test was substituted. For repeated observations, a repeated-measures ANOVA was applied when residuals were normally distributed. If only two repeated levels were compared, a paired-samples *t*-test was used; when the normality assumption was violated a Wilcoxon signed-rank test (two levels) or a Friedman test (≥ 3 levels) replaced the parametric test. Sphericity was assessed with Mauchly’s test, and the Greenhouse–Geisser correction was applied when sphericity was violated. Post-hoc pairwise comparisons after any omnibus ANOVA were adjusted with the Bonferroni procedure. Linear relations between two continuous, normally distributed variables were assessed with Pearson’s *r*; otherwise Spearman’s ρ was used. Rayleigh test was used to assess non-uniformity of spike-phase coupling and parametric Watson-Williams multi-sample test was used for equal means. The threshold for statistical significance was set at two-tailed *α* = 0.05. The fitting and statistical analyses were performed using the MATLAB software package (MathWorks).

### Data availability

The datasets generated and analyzed during this study are available from the corresponding author upon reasonable request.

## Acknowledgments

This manuscript was prepared with the assistance of AI-based tools (ChatGPT and Gemini). ChatGPT was used for improving the clarity, grammar, and overall flow of the text. Both AI-based tools were used for providing guidance on statistical analyses, including verification and refinement of the statistical treatment of results. In all cases, the authors thoroughly reviewed, edited, and validated any AI-generated content. AI-based tools were not used for core research tasks such as experimental design, data acquisition, primary data analysis, interpretation of results, or drawing scientific conclusions. The authors take full responsibility for all content of the manuscript.

## Author contribution

PF designed the study and wrote the manuscript; NE performed experiments and analyses; MC, GL, AA, and AL-V contributed to data acquisition. The authors declare no competing interests.

