## Supplemental figures for "Infraslow modulation of theta synchrony in the hippocampus circuit during REM sleep"

### Supplementary figures

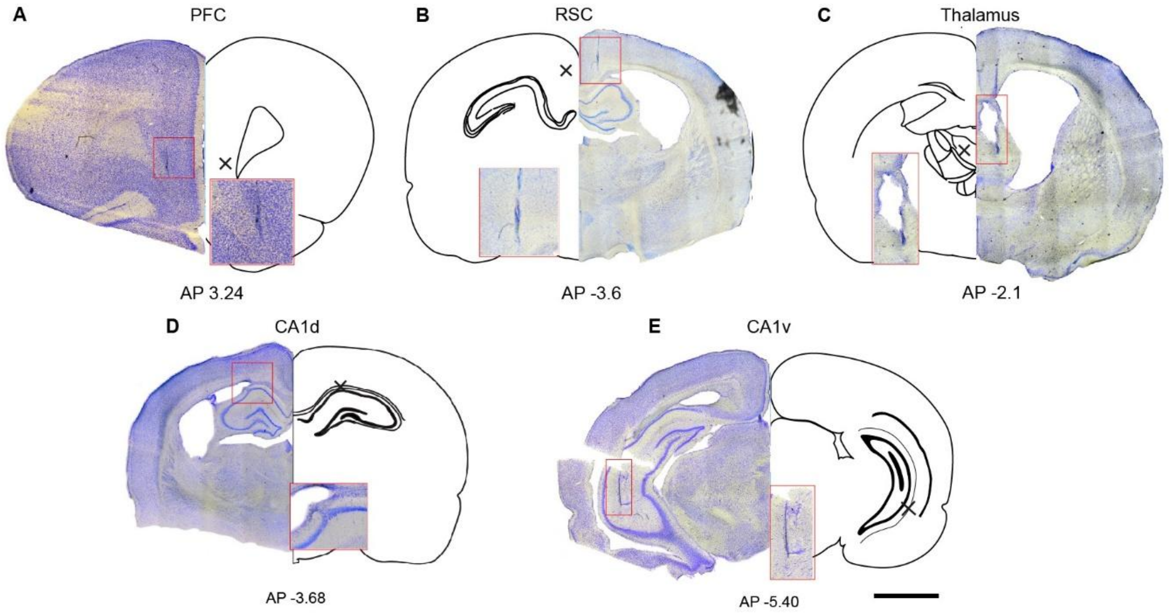


**Supplementary Figure 1. Histological verification of recording sites.** Example Nissl-stained coronal brain sections (rat PF39) illustrating the locations of electrode tracks and recording sites in the prefrontal cortex (PFC) (*A*), retrosplenial cortex (RSC) (*B*), dorsal thalamus (Thalamus) (*C*), dorsal CA1 (CA1d) (*D*), and ventral CA1 (CA1v) (*E*). For each panel, the left image shows the histological section, and the right schematic outlines the corresponding coronal level from a standard rat brain atlas. Red boxes indicate the approximate positions of electrode tips, and insets provide higher-magnification views of the targeted regions. Anterior–posterior (AP) coordinates (in mm relative to bregma) are indicated below each panel. These sections confirm accurate placement of recording electrodes within the intended cortical, hippocampal, and thalamic structures across animals.

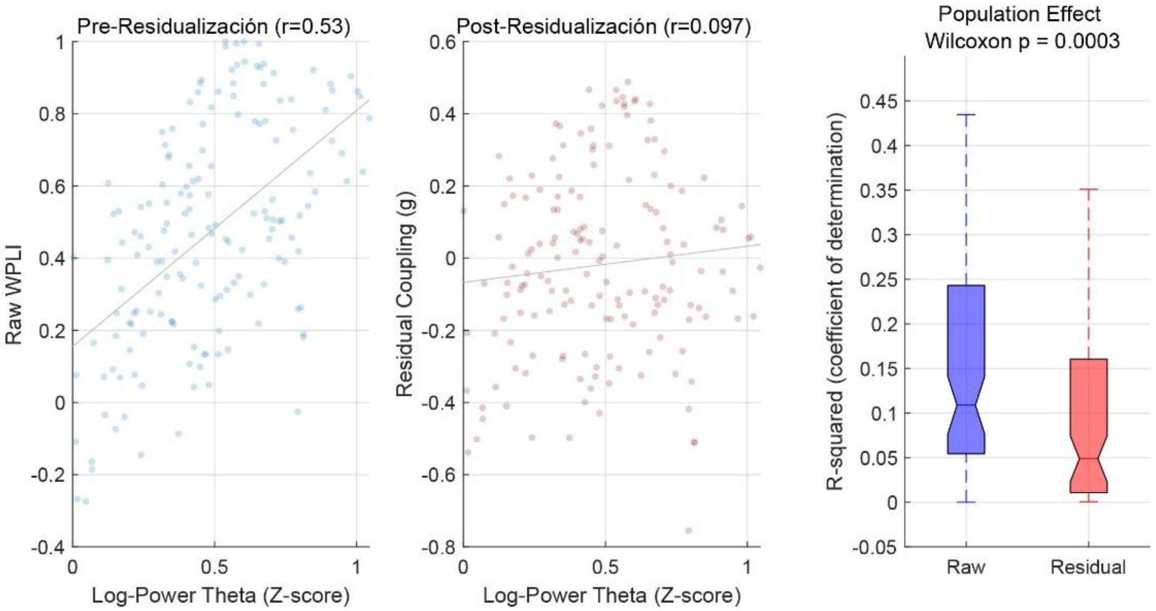


**Supplementary Figure 2. Residualization of theta-band coupling removes power-related confounds.** *Left,* Relationship between raw theta-band weighted Phase Lag Index (wPLI) and log-transformed theta power (z-scored) during REM sleep. Raw wPLI values show a substantial positive correlation with theta power (example session shown; Pearson's r = 0.53), indicating that apparent changes in coupling strength can be partially driven by local signal amplitude. *Middle,* The same relationship after residualization of wPLI with respect to theta power in both signals. Following regression, the correlation between residual coupling (g) and theta power is strongly reduced (r = 0.097), demonstrating effective removal of power-related variance. *Right,* Population summary across animals showing the coefficient of determination ($R^2$) for raw (blue) and residualized (red) coupling. Residualization significantly reduced the proportion of coupling variance explained by theta power (Wilcoxon signed-rank test, P = 0.0003). Together, these analyses confirm that subsequent infraslow fluctuations in coupling reflect genuine changes in interregional phase coordination rather than trivial covariation with local theta power.

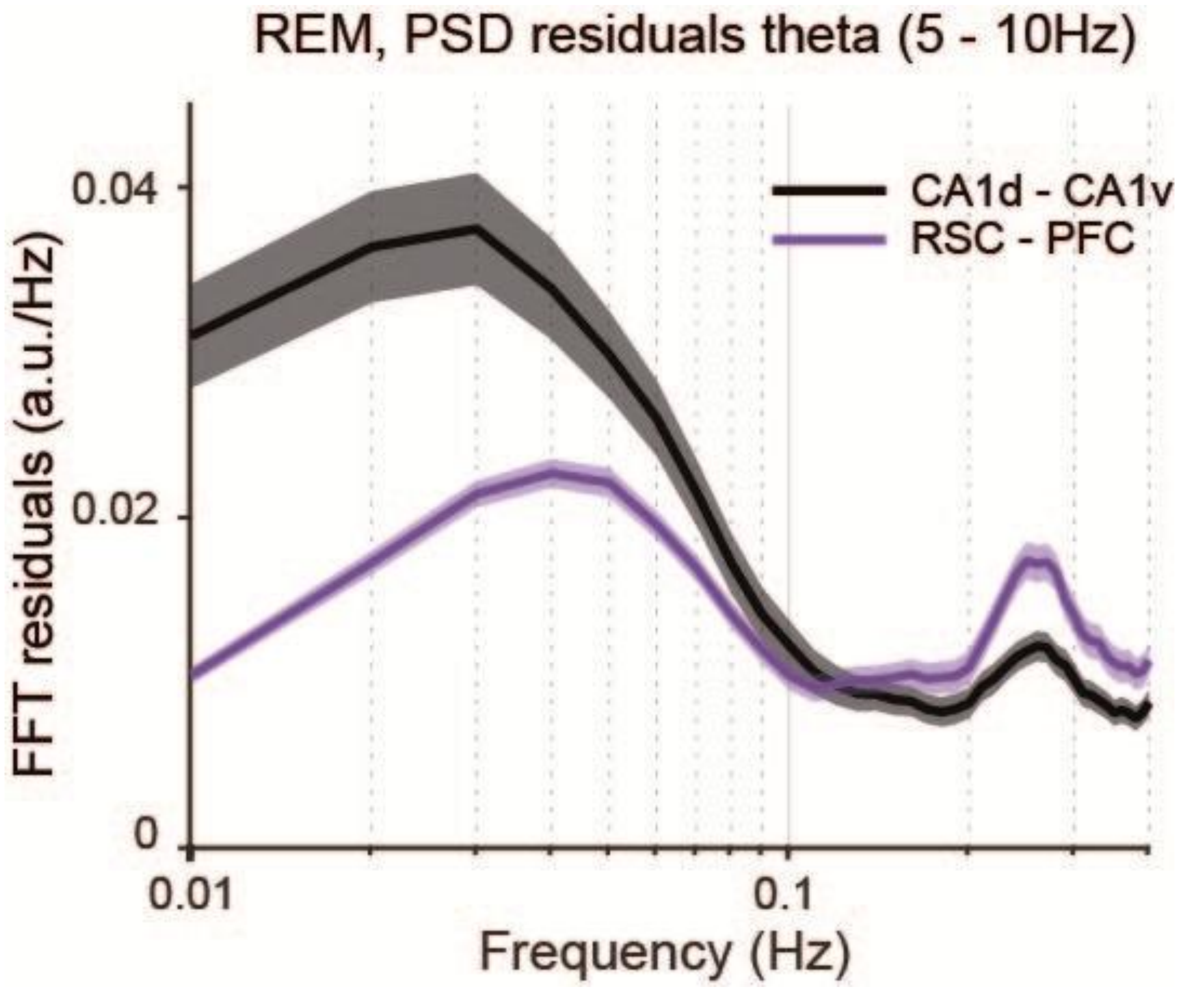


**Supplementary Figure 3. Infraslow modulation of theta-band power residuals during REM sleep.** Power spectral density (PSD) of theta-band (5–10 Hz) power residuals computed during REM sleep for hippocampal (CA1d–CA1v) and cortical (RSC–PFC) signal pairs. Traces show the mean residual spectrum across animals, with shaded areas indicating the standard error of the mean (SEM). After removal of state-specific mean theta power, hippocampal signals exhibited a prominent infraslow peak in the ~0.02–0.04 Hz range, whereas the corresponding cortical residuals displayed a markedly smaller modulation. These results indicate that infraslow dynamics during REM sleep preferentially affect hippocampal theta activity, with substantially weaker expression in cortical networks, supporting the regional specificity of the infraslow modulation observed in the main analyses.

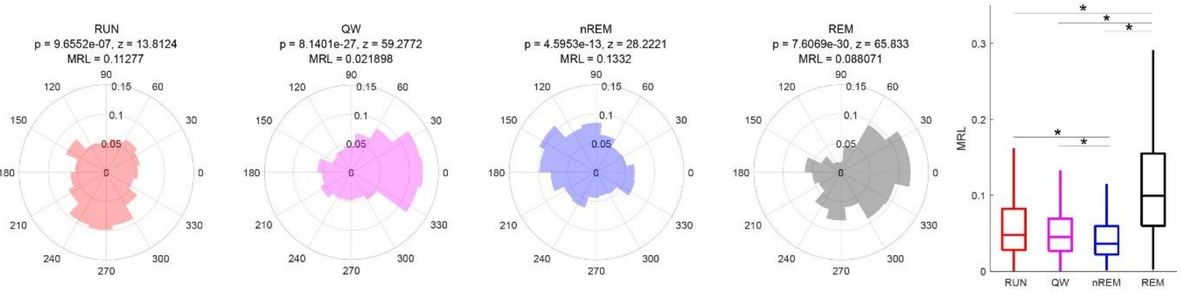


**Supplementary Figure 4. State-dependent phase locking of neuronal firing to the infraslow hippocampal theta coupling signal.** Circular histograms (left) illustrate the distribution of spike phases relative to the infraslow phase of CA1d–CA1v theta synchrony during active exploration (RUN), quiet wakefulness (QW), non–rapid eye movement sleep (nREM), and rapid eye movement sleep (REM). For each state, pooled spikes across regions are plotted, and the resultant vector (black arrow) indicates the preferred phase and strength of phase locking. Rayleigh test statistics (Z and P values) and mean resultant length (MRL) are indicated above each polar plot. Significant non-uniform phase locking is weak or absent during RUN, QW, and nREM, but becomes robust during REM sleep. Right panel summarizes the mean resultant length (MRL) across states. REM sleep exhibits significantly stronger phase locking compared to all other vigilance states, indicating enhanced and consistent entrainment of neuronal firing to the infraslow modulation of hippocampal theta coupling specifically during REM sleep. Box plots show median and interquartile range; whiskers denote data range. Asterisks indicate significant pairwise differences (*$P < 0.05$).
