## Supplemental tables for "Infraslow modulation of theta synchrony in the hippocampus circuit during REM sleep"

**Supplementary Tables**

| Animal | CA1d | CA1v | RSC | PFC | Th |
|---|---|---|---|---|---|
| **AA04** | + | + | + | + | + |
| **AA06** | + | + | + | + | + |
| **AL25** | + | + | + | + | - |
| **MC22** | + | + | + | + | - |
| **MC25** | + | + | + | + | - |
| **NE01** | + | + | + | + | - |
| **PF07** | + | + | + | + | - |
| **PF11** | + | + | + | + | - |
| **PF19** | + | + | + | + | - |
| **PF24** | + | + | + | + | - |
| **PF31** | + | + | + | + | + |
| **PF33** | + | + | + | + | + |
| **PF39** | + | + | + | + | + |
| **PF40** | + | + | + | + | + |
| **TM36** | + | + | + | + | - |

**Table S1**. **Summary of electrode locations across recorded brain areas**. Table shows the availability of viable local field potential recordings for each animal across the targeted brain areas: dorsal and ventral hippocampus (CA1d, CA1v), prefrontal cortex (PFC), retrosplenial cortex (RSC), and dorsal thalamus (Th). A plus sign (+) represents histologically confirmed electrode location and inclusion of the signals in the data analysis, while a minus sign (-) indicates that data from that specific area was unavailable due to missed targeting.

**Analysis of variance**

| Source | Sum Sq. | d.f. | Mean Sq. | F | Prob > F | |
|---|---|---|---|---|---|---|
| **area** | 0.02734316 | 1 | 0.02734316 | 71.0025169 | 1.30 x$10^{-16}$ | (*) |
| **state** | 0.20906443 | 3 | 0.06968814 | 180.960576 | 7.44 x$10^{-93}$ | (*) |
| **area*state** | 0.00991082 | 3 | 0.00330361 | 8.57854435 | 1.27 x$10^{-5}$ | (*) |
| **Error** | 0.36700148 | 953 | 0.0003851 | | | |
| **Total** | 0.61175838 | 960 | | | | |

**Pairwise multiple comparison (Bonferroni)**

| state | area1 | area2 | P | |
|---|---|---|---|---|
| RUN | CA1d | CA1v | 1.51 x$10^{-7}$ | (*) |
| QW | CA1d | CA1v | 0.116829 | n.s. |
| nREM | CA1d | CA1v | 0.979659 | n.s. |
| REM | CA1d | CA1v | 5.99 x$10^{-8}$ | (*) |

**Table S2. Statistical analysis of delta band power across brain states.** Summary of the two-way ANOVA performed to evaluate the effects of brain area (CA1d vs. CA1v) and sleep-wake state (RUN, QW, nREM, REM) on delta oscillation power (0.5–2 Hz). The table presents the sum of squares, degrees of freedom (d.f.), F-statistics, and P-values for main effects and interaction. Pairwise multiple comparisons (Bonferroni corrected) are shown below, highlighting significant regional differences particularly during activated states.

**Analysis of variance**

| Source | Sum Sq. | d.f. | Mean Sq. | F | Prob > F | |
|---|---|---|---|---|---|---|
| **area** | 0.31456099 | 1 | 0.31456099 | 298.479127 | $2.18 \times 10^{-58}$ | (*) |
| **state** | 0.96003234 | 3 | 0.32001078 | 303.650299 | $2.59 \times 10^{-138}$ | (*) |
| **area*state** | 0.20054162 | 3 | 0.06684721 | 63.4296592 | $2.14 \times 10^{-37}$ | (*) |
| **Error** | 1.00434702 | 953 | 0.00105388 | | | |
| **Total** | 2.47987284 | 960 | | | | |

**Pairwise multiple comparison (Bonferroni)**

| state | area1 | area2 | p-val | |
|---|---|---|---|---|
| RUN | CA1d | CA1v | $1.51 \times 10^{-7}$ | (*) |
| QW | CA1d | CA1v | $2.09 \times 10^{-4}$ | (*) |
| nREM | CA1d | CA1v | 0.999996 | n.s. |
| REM | CA1d | CA1v | $5.99 \times 10^{-8}$ | (*) |

**Table S3. Statistical analysis of theta band power across brain states.** Summary of the two-way ANOVA assessing the influence of brain area and state on theta oscillation power (5–10 Hz). The analysis reveals significant main effects for both area and state, as well as significant interaction. Post-hoc Bonferroni comparisons indicate that theta power differences between dorsal and ventral hippocampus are most pronounced during RUN and REM sleep.

**Analysis of variance**

| Source | Sum Sq. | d.f. | Mean Sq. | F | Prob > F | |
|---|---|---|---|---|---|---|
| **area** | 1997.02766 | 1 | 1997.02766 | 171.8938106 | $3.17 \times 10^{-36}$ | (*) |
| **state** | 4810.430914 | 3 | 1603.476971 | 138.0190032 | $2.94 \times 10^{-74}$ | (*) |
| **area*state** | 1815.70334 | 3 | 605.2344467 | 52.09545039 | $3.50 \times 10^{-31}$ | (*) |
| **Error** | 11071.762 | 953 | 11.6177985 | | | |
| **Total** | 19439.6213 | 960 | | | | |

**Pairwise multiple comparison (Bonferroni)**

| state | area1 | area2 | p-val | |
|---|---|---|---|---|
| RUN | CA1d | CA1v | $5.99 \times 10^{-8}$ | (*) |
| QW | CA1d | CA1v | 0.855398 | n.s. |
| nREM | CA1d | CA1v | 1 | n.s. |
| REM | CA1d | CA1v | $5.99 \times 10^{-8}$ | (*) |

**Table S4. Statistical analysis of the theta/delta power ratio.** Summary of the two-way ANOVA for the theta/delta power ratio, calculated to quantify the spectral shift between states. Results show significant effects for area, state, and their interaction, confirming that the dominance of theta over delta activity is state-dependent and differentially expressed along the septotemporal axis.

**Analysis of variance**

| Source | Sum Sq. | d.f. | Mean Sq. | F | Prob > F |
|---|---|---|---|---|---|
| **area** | 1105.4 | 1 | 1105.36 | 30.8 | 0 |
| **state** | 920.5 | 3 | 306.83 | 8.55 | 0 |
| **area*state** | 273.2 | 3 | 91.08 | 2.54 | 0.0552 |
| **Error** | 48095.6 | 1340 | 35.89 | | |
| **Total** | 50190.8 | 1347 | | | |

**Pairwise multiple comparison (Bonferroni)**

| state1 | state2 | p-val | |
|---|---|---|---|
| RUN | QW | 0.0252 | (*) |
| RUN | nREM | 0.415 | n.s. |
| RUN | REM | 0.00574 | (*) |
| QW | nREM | 0.16 | n.s. |
| QW | REM | 0.00000181 | (*) |
| nREM | REM | 0.000511 | (*) |

**Table S5. Statistical analysis of mean firing rate (MUA).** Summary of the two-way ANOVA investigating variations in multi-unit activity (MUA) across hippocampal regions and brain states. The table details the statistical parameters for main effects and the interaction term. Pairwise comparisons (Bonferroni adjusted) show that firing rates are significantly modulated by state, reaching highest levels during REM sleep compared to QW and nREM.

**Watson-Williams multi-sample test**

| | **CA1d** | **CA1v** | **RSC** | **PFC** | **Th** |
|---|---|---|---|---|---|
| **CA1d** | - | $1.2x10^{-5}$ (*) | 0.24 | 0.045 (*) | $7.7x10^{-12}$ (*) |
| **CA1v** | | - | 0.004 (*) | 0.01 (*) | 0.24 |
| **RSC** | | | - | 0.55 | $7.3x10^{-6}$ (*) |
| **PFC** | | | | - | $9.4x10^{-6}$ (*) |
| **Th** | | | | | - |

**Table S6. Pairwise comparisons of spike-phase locking preferences (Watson-Williams test).** Results of the Watson-Williams multi-sample test comparing the mean phase angles of neuronal spiking relative to the infraslow theta coupling cycle across recorded regions. The table displays P-values for pairwise comparisons between regions (CA1d, CA1v, RSC, PFC, Th), indicating significant differences in the preferred phase of entrainment among specific anatomical structures during REM sleep.
